# The limits of long-term selection against Neandertal introgression

**DOI:** 10.1101/362566

**Authors:** Martin Petr, Svante Pääbo, Janet Kelso, Benjamin Vernot

## Abstract

Several studies have suggested that introgressed Neandertal DNA was subjected to negative selection in modern humans due to deleterious alleles that had accumulated in the Neandertals after they split from the modern human lineage. A striking observation in support of this is an apparent monotonic decline in Neandertal ancestry observed in modern humans in Europe over the past 45 thousand years. Here we show that this apparent decline is an artifact caused by gene flow between West Eurasians and Africans, which is not taken into account by statistics previously used to estimate Neandertal ancestry. When applying a more robust statistic that takes advantage of two high-coverage Neandertal genomes, we find no evidence for a change in Neandertal ancestry in Western Europe over the past 45 thousand years. We use whole-genome simulations of selection and introgression to investigate a wide range of model parameters, and find that negative selection is not expected to cause a significant long-term decline in genome-wide Neandertal ancestry. Nevertheless, these models recapitulate previously observed signals of selection against Neandertal alleles, in particular a depletion of Neandertal ancestry in conserved genomic regions that are likely to be of functional importance. Thus, we find that negative selection against Neandertal ancestry has not played as strong a role in recent human evolution as had previously been assumed.

## Introduction

Interbreeding between Neandertals and modern humans approximately 55,000 years ago has resulted in all present-day non-Africans inheriting at least 1-2% of their genomes from Neandertal ancestors (1, 2). There is significant heterogeneity in the distribution of this Neandertal DNA across the genomes of present-day people (3, 4), including a reduction in Neandertal alleles in functionally constrained genomic regions (3). This has been interpreted as evidence that some Neandertal alleles were deleterious for modern humans and were subject to negative selection following introgression (3, 5). Several studies have suggested that low effective population size (*N_e_*) in Neandertals led to decreased efficacy of purifying selection and accumulation of weakly deleterious variants. Following introgression, these deleterious alleles, along with linked neutral Neandertal alleles, would have been subjected to more efficient purifying selection in the modern human population (6, 7).

In apparent agreement with this hypothesis, a study of Neandertal ancestry in a set of anatomically modern humans from Upper-Paleolithic Europe used two independent statistics to conclude that the amount of Neandertal DNA in modern human genomes decreased monotonically over the last 45 thousand years (8). This decline was interpreted as direct evidence for continuous negative selection against Neandertal alleles in modern humans (8–11). However, it was not formally shown that selection on deleterious introgressed variants could produce a decline in Neandertal ancestry of the observed magnitude. Nevertheless, the decreasing Neandertal ancestry in modern humans over time, together with the suggestion of a higher burden of deleterious alleles in Neandertals are now commonly invoked to explain the fate of Neandertal ancestry in modern humans (9–12).

Here, we re-examine estimates of Neandertal ancestry in ancient and present-day modern humans, taking advantage of a second high-coverage Neandertal genome that recently became available (13). This allows us to avoid some key assumptions about modern human demography that were made in previous studies. Our analysis shows that the Neandertal ancestry proportion in non-Africans has not decreased significantly through the last 45,000 years. We then compare this result to realistic, genome-scale simulations, and confirm that under a model of weak selection against Neandertal alleles, after an initial sharp decrease in the Neandertal ancestry in modern humans, an indefinite period during which this ancestry remains constant in the population is expected.

## Results

### Previous Neandertal ancestry estimate

A number of methods have been developed to quantify Neandertal ancestry in modern human genomes (14). Among the most widely used is the *f_4_*-ratio statistic, which measures the fraction of drift shared with one of two parental lineages to determine the proportion of ancestry *α* contributed by that lineage (Fig. 1C and 1D) (15, 16). Although it has been used to draw inferences about gene-flow between archaic and modern human populations, *f_4_*-ratio statistics are known to be sensitive to violations of the underlying population model (15). Estimating *α*, the proportion of ancestry contributed by lineage *A* to *X*, requires a sister lineage *B* to lineage *A* which does not share drift with *X* after separation of *B* from *A* (Fig. 1C, 1D). Fu *et al*. used an *f_4_*-ratio statistic to infer the contribution from an archaic lineage by first estimating the proportion of East African ancestry in a non-African individual *X*, under the assumption that Central and West Africans (*A*) are an outgroup to the East African lineage (*B*) and to the modern human ancestry in non-Africans (8). The proportion of Neandertal ancestry was then calculated simply as *1 – α*, under the assumption that all ancestry that is not of East African origin must come from an archaic lineage (Fig. 1C). We refer to this statistic as an “indirect *f*_*4*_-ratio”.

**Figure 1.**
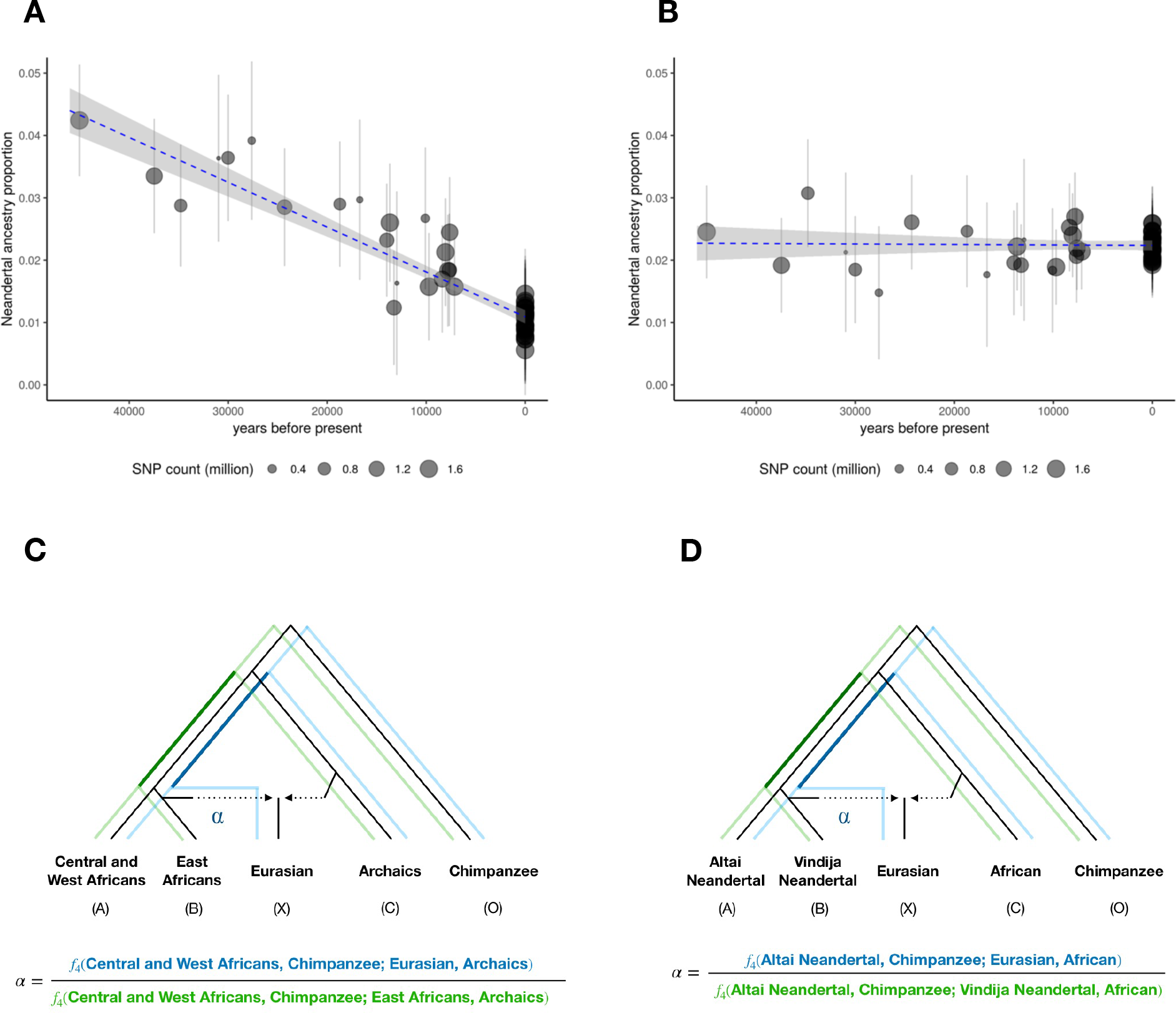
Direct and indirect *f_4_*-ratio estimates of Neandertal ancestry. **A)** Indirect and **B)** Direct *f_4_*-ratio estimates of Neandertal ancestry in ancient and modern West Eurasians (black points), and best fit line, weighted by number of sites genotyped in each sample (dotted line, shaded area is 95% CI). **C)** Tree model used for indirect *f_4_*-ratio. **D)** Tree model used for direct *f_4_* ratio, utilizing two high coverage Neandertal genomes. Green and blue lines in C) and D) indicate drift paths taken through each population tree in the numerator (blue) and denominator (green) of the overall *f_4_*-ratio statistic. Present-day individuals are Europeans from the SGDP panel (29).

Given the sensitivity of the *f_4_*-ratio method to violations of the underlying population models (15), we explored the validity of assumptions on which this calculation was based. In addition to the topology of the demographic tree, the indirect *f_4_*-ratio assumes that the relationship between Africans and West Eurasians has remained constant over time (8).

To test for gene-flow between Africans and West Eurasians over time, we calculated D statistics in the form of *D(Ust’-Ishim, X; African population, Chimp)*, which tests for changes in affinity (i.e. the number of shared derived alleles) between a series of West Eurasians (*X*) and Africans with respect to Ust’-Ishim - a 45,000 year-old Eurasian individual who is the oldest modern human genome in this dataset. In the absence of gene-flow between a West Eurasian lineage *X* and Africans in this time-frame, the value of this *D* statistic will not be significantly different from 0 for any of the West Eurasians regardless of their age. However, in violation of the assumption of the indirect *f_4_*-ratio, we find that starting from as early as 20 thousand years ago (kya) this *D* statistic becomes increasingly negative, consistent with gene-flow between West Eurasians and Africans during this time (Fig. 2). To rule out potential technical issues, we repeated this analysis substituting Oceanians for Africans and find no changes in affinity over time (Fig. 2). We note that while the increase in affinity between African and West Eurasian populations shows that the assumption of a constant relationship between these populations over time is incorrect, it is not informative about the direction of the gene-flow.

**Figure 2.**
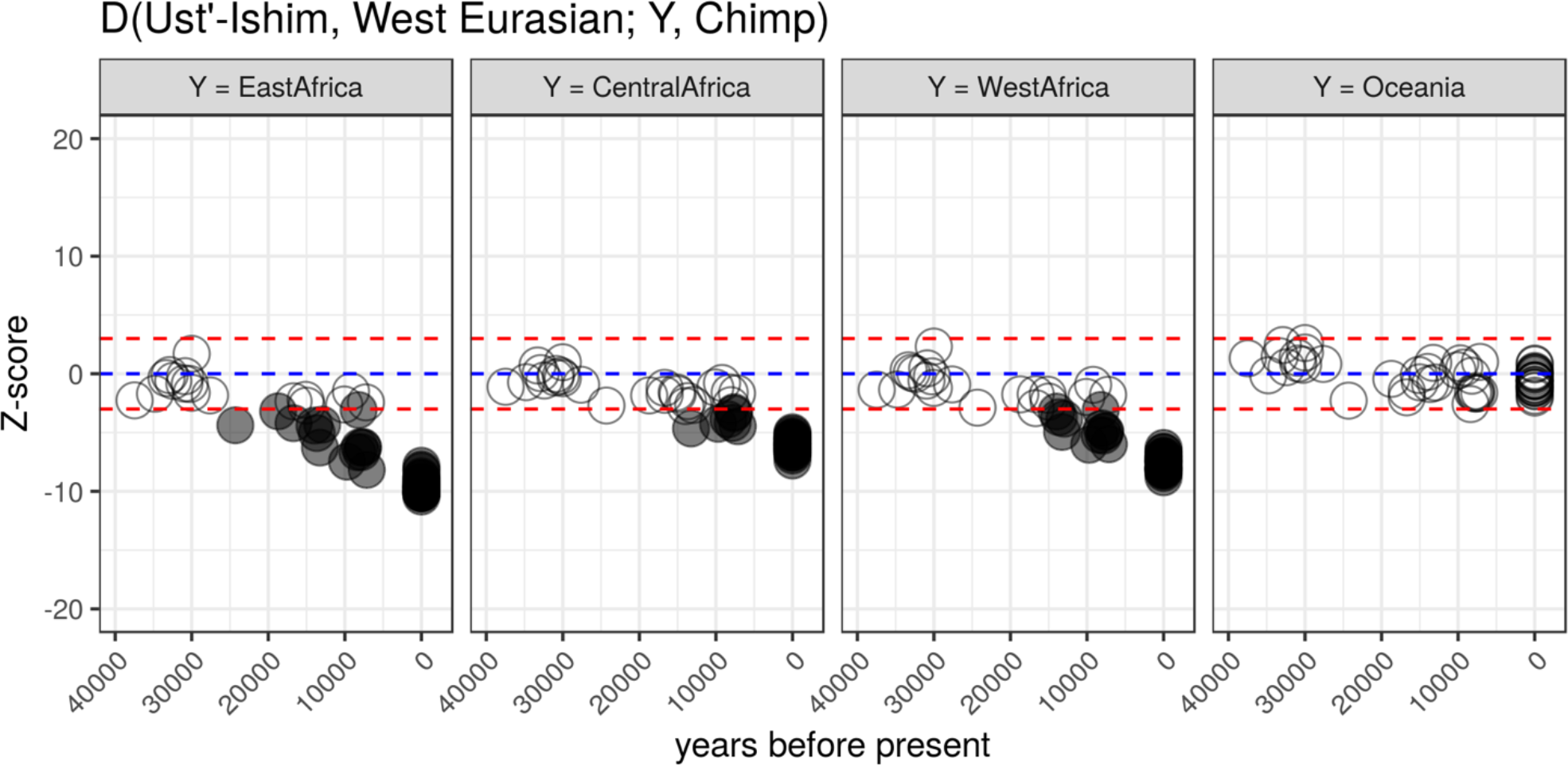
Affinity of ancient and modern West Eurasian individuals to three African populations (left) and Oceanians (right). Z score values for *D(Ust’-Ishim, X; African population or Oceanians, Chimp)*, where X is a series of European individuals over time. Negative Z-scores indicate that an individual X is significantly closer to a population in the third position of the *D* statistic than Ust’-Ishim is. Assuming all non-Africans share a common ancestor after the split from an African lineage, this can be interpreted as evidence of gene-flow between Europeans and Africans over time and no such gene flow between Europeans and Oceanians.

### A robust statistic to estimate Neandertal ancestry

The recent availability of a second high-coverage Neandertal genome allows us to estimate Neandertal ancestry using two Neandertals – an individual from the Altai Mountains, the so-called “Altai Neandertal” (17) and an individual from Vindija Cave in Croatia, the so-called “Vindija Neandertal” (13). Specifically, we estimate the proportion of ancestry coming from the Vindija lineage into a modern human (*X*) using the Altai Neandertal as a second Neandertal in an *f_4_*-ratio calculated as *f_4_(Altai, Chimp; X, African) / f4(Altai, Chimp; Vindija, African)* (Fig. 1D), which we refer to as a “direct *f_4_*-ratio”. Note that unlike the indirect *f_4_*-ratio, the *f_4_*-ratio in this formulation does not make assumptions about deep relationships between modern human populations (Fig. 1D). Instead, it assumes that any Neandertal population that contributed ancestry to West Eurasians formed a clade with the Vindija Neandertal population. Recent analyses showed that this is the case for all non-African populations studied to date, including the ancient modern humans included in this study (13, 18). Crucially, when we use the direct *f_4_*-ratio to estimate the trajectory of Neandertal ancestry in West Eurasians we observe nearly constant levels of Neandertal ancestry over time (Fig. 1B) and find that a null model of no slope can no longer be rejected (Fig. 1B, p = 0.48).

This result has several implications for our understanding of the fate of Neandertal ancestry in modern humans. First, it constrains the timescale during which selection could have significantly affected the average genome-wide Neandertal ancestry in modern humans, an issue addressed below in more detail. Second, it has consequences for the so-called “dilution” hypothesis, which suggests that the larger proportion of Neandertal ancestry in East Eurasians compared to West Eurasians can be explained by West Eurasian admixture with a “Basal Eurasian” population who may have carried less Neandertal ancestry than other non-Africans (19, 20). We find that a statistic informative for Basal Eurasian ancestry (*f_4_(West Eurasian W, East Asian X; Ust’-Ishim, Chimp)*), is significantly negative in present-day West Eurasians (Fig. S1), implying that all West Eurasians have a contribution from a lineage that predates the split of East and West Eurasians. However, our finding that there is no significant decline in Neandertal ancestry in West Eurasians suggests that admixture with this population did not influence their Neandertal ancestry in a significant way (Fig. 1B). This is consistent with a previous analysis of an early European farmer from Germany, who was found to have carried a similar amount of Neandertal ancestry to present-day Europeans (~1.8%), despite having a significant proportion of her genome derived from Basal Eurasian ancestry (44%) (21).

### Testing the robustness of Neandertal ancestry statistics

Gene-flow between West Eurasians and Africans is evident in our *D* statistic results (Fig.2), and has also been detected by earlier studies of present-day Africans (22, 23). Crucially, such gene-flow violates the assumptions of the indirect *f_4_*-ratio, but it is not clear exactly how this statistic will be affected. Furthermore, the direction of gene-flow may be of interest, yet cannot be determined by the *D*-statistics presented above. Thus, we simulated several scenarios of migration between West Eurasians and Africans post-Neandertal introgression, and calculated both direct and indirect *f4*-ratios on these simulated sequences, along with the true Neandertal proportion (Fig. 3).

**Figure 3.**
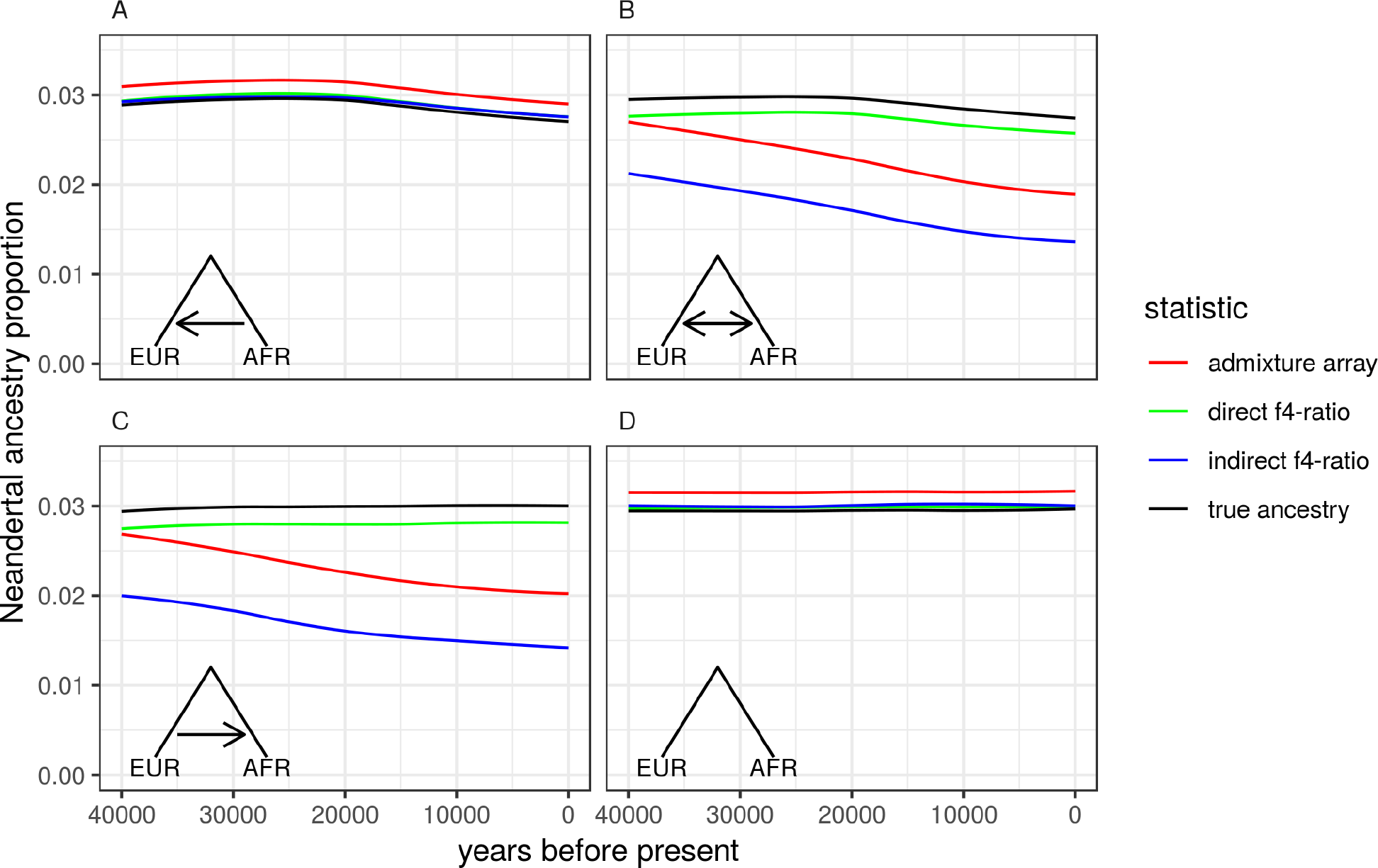
Neandertal ancestry estimates in neutral simulations of migration. True Neandertal ancestry proportions as a function of time (black solid lines) and estimates of Neandertal ancestry using three different statistics on simulated genomic data. Genomic data was simulated under four models of migration. **A)** Migration from Africans to West Eurasians, at a rate of 0.0001 migrants per generation, beginning 20 kya; **B)** bi-directional migration between West Eurasia and Africa; **C)** migration from West Eurasians to Africans, using the same parameters as (A); **D)** no migration.

In short, we find that even moderate levels of gene flow from West Eurasians into Africans (0.0001 migrants per generation, starting at 20 kya) lead to mis-estimates of Neandertal ancestry when using the indirect *f_4_*-ratio statistic (Fig. 3C). Such scenarios result in depressed estimates of Neandertal ancestry, with this effect being more pronounced in simulated genomes sampled closer to the present day, incorrectly resulting in an apparent decline in Neandertal ancestry over time. This is qualitatively consistent with the apparent decline in Neandertal ancestry when using the indirect *f*_*4*_-ratio on real West Eurasian individuals, suggesting that the previous observations are artifacts produced by gene flow from West Eurasia into Africa.

In contrast, migration only from Africa to West Eurasia results in a true decline of Neandertal ancestry, due to its replacement by modern human alleles. In this scenario, the true decline is accurately estimated by both the indirect and direct *f_4_*-ratios (Fig. 3A). In scenarios of bi-directional migration, this true decline is only accurately measured by the direct *f_4_*-ratio (Fig. 3B).

An independent statistic, using a different set of genomic sites in the same ancient individuals, has been used as a second line of evidence for an ongoing decrease in Neandertal ancestry (8). This statistic, which we refer to as the “admixture array statistic”, measures the proportion of Neandertal-like alleles in a given sample at sites where present-day Yoruba individuals carry a nearly-fixed allele that differs from homozygous sites in the Altai Neandertal (24). Using the simulations discussed above, we selected genomic sites using the same conditioning. We then calculated the proportion of Neandertal-like alleles in each simulated West Eurasian genome. Much like the indirect *f*_*4*_-statistic, the admixture array statistic is affected by gene flow from West Eurasians into Africans, incorrectly inferring a decline of Neandertal ancestry (Fig. 3C).

We note that the direct *f_4_*-ratio is not immune to the effects of migration from West Eurasia to Africa. Specifically, the direct *f_4_*-ratio measures the amount of Neandertal ancestry in West Eurasians in excess of any Neandertal ancestry present in Africans. Thus, it is likely that even our updated estimates of Neandertal ancestry are in fact underestimates.

### Long-term dynamics of selection against introgressed DNA

Our observation that Neandertal ancestry levels did not significantly decrease from about 40,000 years ago until today is seemingly at odds with the hypothesis that lower effective population sizes in Neandertals led to an accumulation of deleterious alleles, which were then subjected to negative selection in modern humans (3, 8–10). To investigate the expected long-term dynamics of selection against Neandertal introgression under this hypothesis, we simulated a model of the human genome with empirical distributions of functional regions and selection coefficients, extending a strategy previously applied by Harris and Nielsen (6). We simulated modern human and Neandertal demography, including a low long-term effective population size (*N_e_*) in Neandertals (Neandertal *N_e_* = 1,000 vs modern human *N_e_* = 10,000) and 10% introgression at 55 kya (2200 generations ago, assuming generation time of 25 years). To track the changes in Neandertal ancestry following introgression, we placed fixed Neandertal-human differences as neutral markers, both between regions that accumulated deleterious mutations (to study the effect of negative selection on linked genome-wide neutral Neandertal variation) as well as within regions directly under selection (to track the effect of negative selection itself) (Fig. 4A).

**Figure 4.**
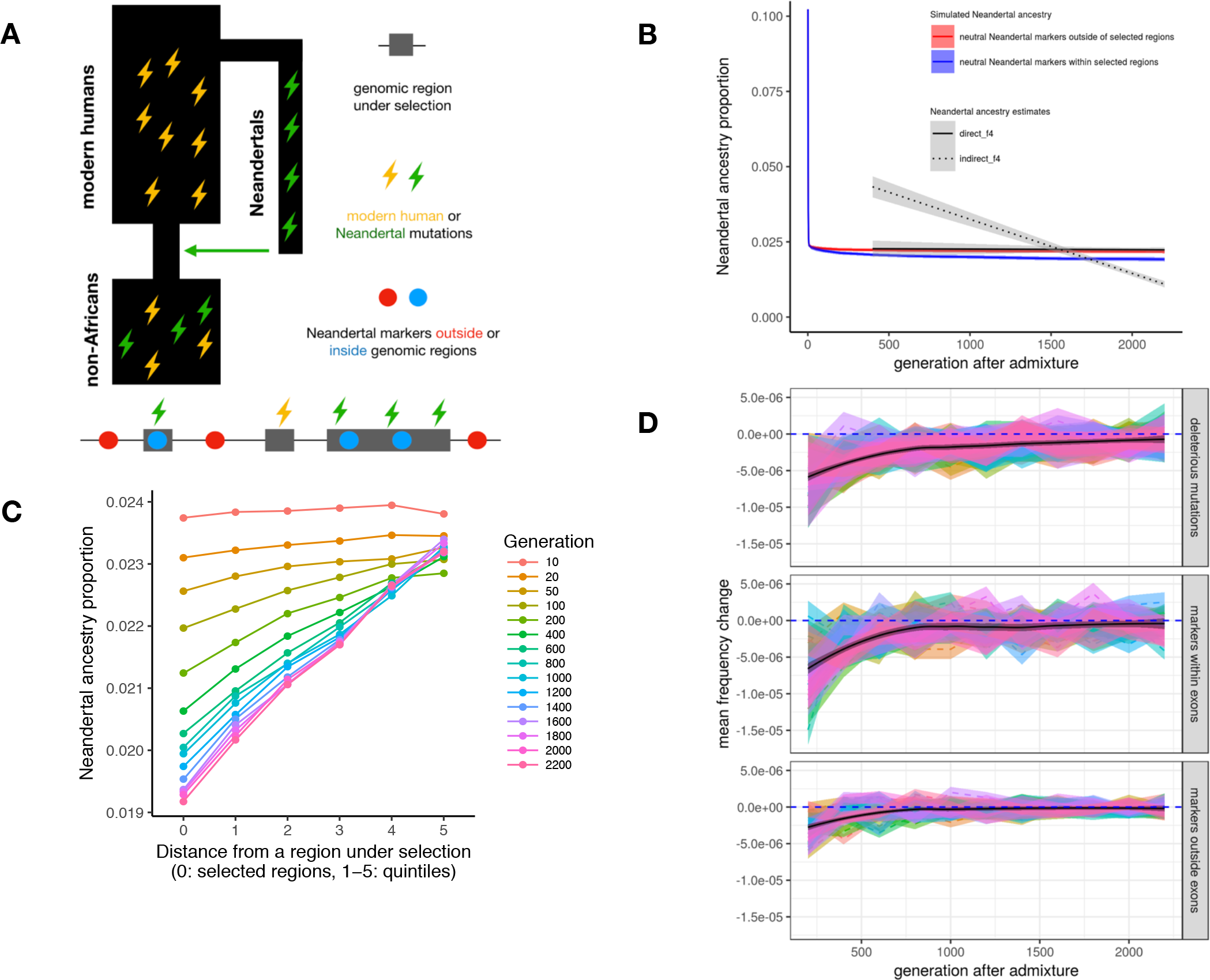
Simulations of selection against Neandertal ancestry. **A)** Schematic of simulated selection in Neandertals and modern humans. Deleterious mutations (lightning bolts) accumulate in 70 Mb of realistically distributed exonic sequence, in both modern humans and Neandertals. Neutral Neandertal “ancestry informative” markers are placed within (blue dots) and between (red dots) functional regions under selection to track the presence of Neandertal ancestry along a chromosome. Results from simulations with different amounts of deleterious sequence are shown in Fig. S6. **B)** Simulated Neandertal ancestry proportions across 55 ky, at neutral markers within (blue) and between (red) selected regions. Empirical observations from Figs. 1B and 2B are shown in black lines, for comparison. **C)** Depletion of simulated Neandertal ancestry at neutral markers as a function of distance to selected regions (bins 0 to 5: markers in bin 0 are those falling within selected regions – blue dots in panel A; bins 1-5 represent quintiles of neutral markers between selected regions). Depletion is stronger closer to selected regions, and this effect increases over time. Initial introgression levels are simulated at 10%. **D)** Changes in frequencies of fixed neutral Neandertal markers and deleterious Neandertal exonic mutations over time, starting from generation 200 – using the same simulation data as in the plot in panel B.

Similar to Harris et al., we observed abrupt removal of Neandertal alleles from the modern human population during the first ~10 generations after introgression, followed by quick stabilization of Neandertal ancestry levels (Fig 4B). When compared with empirical estimates of Neandertal ancestry, we find a better fit between these simulations and the direct *f_4_*-ratio estimate than with the indirect *f_4_*-ratio estimate, suggesting that our direct Neandertal ancestry estimates are consistent with theoretical expectations of genome-wide selection against introgression (Fig 4B; root mean squared error (RMSE) between mean simulated Neandertal ancestry estimates and direct *f_4_*-ratio estimates = 0.0005, RMSE between simulations and indirect *f_4_*-ratio estimates = 0.01).

Because many factors can potentially influence the efficacy of negative selection, and no model fully captures all of these, we next sought to determine whether there is a combination of model parameters that could potentially lead to long-term continuous removal of Neandertal ancestry over time. Surprisingly, we failed to find a model which would produce a significant decline over time, although we tried: (i) decreasing the long-term Neandertal *N_e_* prior to introgression (making purifying selection in Neandertals even less efficient), (ii) increasing the N_e_ of modern humans after introgression (i.e. increasing the efficacy of selection against introgressed alleles), (iii) artificially increasing the deleteriousness of Neandertal variants after introgression (approximating a “hybrid incompatibility” scenario), or (iv) simulating mixtures of dominance coefficients (Fig. S3-S7). Varying these factors primarily affected the magnitude of the initial removal of introgressed DNA by increasing the number of perfectly linked deleterious mutations in early Neandertal-modern human offspring, which in turn influenced the final level of Neandertal ancestry in the population (Fig. S3-S7).

The depletion of Neandertal ancestry around functional genomic elements in modern human genomes has also been taken as evidence for selection against Neandertal introgressed DNA (3, 8). We next examined the genomic distribution of Neandertal markers at different time-points in our simulations to determine whether our models can recapitulate these signals. In agreement with empirical results in present-day humans (3), we found a strong negative correlation between the proportion of Neandertal introgression surviving at a locus and distance to the nearest region under selection (Fig. 4C). Furthermore, we found that the strength of this correlation increases over time, with the bulk of these changes occurring between 10 and 400 generations post-admixture (mean Pearson’s correlation coefficient ρ = 0.026, 0.8, 0.96 at generations 10, 400 and 2200, respectively (Fig. S9)). We note that this time period predates all existing ancient modern human sequences, frustrating any current comparison to empirical data. Despite no apparent change in average Neandertal ancestry proportion, we observe a smaller though still significant decrease in linked Neandertal ancestry during the time period for which modern human sequences exist (approximately 400-2200 generations post-admixture) (Fig. 4C, 4B). Indeed, by looking at the average per-generation changes in frequencies of simulated Neandertal mutations (that is, derivatives of allele frequencies in each generation), we observe the impact of negative selection on linked neutral Neandertal markers until at least ~700 generations post admixture (Fig. 4D), and find that it closely follows the pattern of frequency derivatives of introgressed deleterious mutations (Fig. 4D). After this period of gradual removal, selection against linked neutral variation slows down significantly as genome-wide Neandertal ancestry becomes largely unlinked from regions that are under negative selection (Fig. 4D). In contrast, the selected variants themselves are still removed, although at increasingly slower rates (Fig. 4D). Due to this slow rate, and the small contribution these alleles make to the overall levels of Neandertal ancestry, their continued removal has little impact on the slope of Neandertal ancestry over time.

### Conclusions

Our re-evaluation of Neandertal ancestry in modern human genomes indicates that overall levels of Neandertal ancestry in West Eurasia have not significantly decreased over the past 45 thousand years, and that previous observations of continuous Neandertal ancestry decline were likely an artifact of unaccounted-for gene-flow from West Eurasia into Africa. Furthermore, we find that negative selection against introgression is expected to have the strongest impact on genome-wide Neandertal ancestry during the first few hundred generations, and becomes negligible in the time frame for which ancient samples are currently available.

Our findings can be extrapolated to other cases where one species or population contributes a fraction of ancestry to another species or population, a frequent occurrence in nature (5, 25–28). Even in cases where the introgressing population carries a high burden of deleterious mutations, negative selection is not expected to result in an extended decrease in the overall genome-wide ancestry contributed by that population. Therefore, any long-term shifts in overall ancestry proportions over time in such situations are likely to be the result of forces other than negative selection, for example admixture with one or more other populations.

## Materials and Methods

### Source code and computational notebooks

Complete source code for data processing and simulation pipelines, as well as R and Python Jupyter notebooks with all analyses, can be downloaded from the project repository on GitHub: https://www.github.com/bodkan/nea-over-time.

### Data processing

SNP data captured at ~2.2 million loci (combination of SNP panels 1, 2, 3, described in (24)) from Upper-Paleolithic individuals published by Fu et al. (8) were obtained in EIGENSTRAT format from the David Reich lab (https://reich.hms.harvard.edu/datasets). Altai and Vindija Neandertal genotypes were converted from VCF to EIGENSTRAT format after filtering the data using the Map35_100% criteria described in (17). For comparisons with present-day populations, we used genotype calls published by the Simons Genome Diversity Project (SGDP) (29), and converted them to EIGENSTRAT format. All data were then combined into single EIGENSTRAT dataset using the “mergeit” command from the ADMIXTOOLS package (15).

SNP data captured using the “archaic admixture array” (described as SNP panel 4 in (24)) published by Fu et al. (8) were also downloaded from the Reich lab website. To enrich for sites that truly originated in the Neandertals, we further restricted to sites homozygous in the Altai and Vindija Neandertal genomes, which results in a set of approximately 480 thousand sites carrying such nearly fixed Yoruba-Neandertal differences.

### Admixture statistics

All D statistics, f_4_ statistics and f_4_-ratio statistics were calculated on the merged 2.2 million loci EIGENSTRAT dataset using our R package admixr (available from https://www.github.com/bodkan/admixr, release v0.1) which utilizes the ADMIXTOOLS software suite for all underlying calculations (15).

### Indirect *f*_*4*_-ratio Neandertal ancestry estimate

Indirect *f_4_*-ratio Neandertal ancestry estimates (Fig. 1A) were calculated on all Upper-Paleolithic humans in our data set, as well as all European individuals from the SGDP human diversity panel. For each individual X in the dataset, the proportion of Neandertal ancestry was calculated as *1 - f*_4_*(West and Central Africans, Chimp; X, Archaics) / f*_4_*(West and Central Africans, Chimp; East African, Archaics)*, where *West and Central Africans* are Yoruba, Mbuti and Mende from the SGDP panel, *East Africans* are SGDP Dinka, and *Archaics* are the Altai Neandertal and the high-coverage Denisovan (30) individuals. This is the f_4_-ratio calculation used in the original Fu et al. study (8).

### Direct *f*_*4*_-ratio Neandertal ancestry estimate

Direct *f_4_*-ratio estimates (Fig. 1B) were calculated on the same data as indirect f_4_-ratio estimates (see above), but using the following setup: *f_4_(Altai, Chimp; X, African) / f_4_(Altai, Chimp; Vindija, African)* for a combined set of African populations Yoruba, Dinka and Mbuti.

### Admixture array proportion

Neandertal ancestry levels using the using the set of nearly-fixed African-Neandertal sites were calculated as a proportion of alleles in a test individual matching the allele seen in the Neandertals.

### Affinity of EMH individuals towards present-day populations over time

We calculated a *D* statistic in the form *D(Ust-Ishim, X; Y, Chimp)*, which tests for changes in the sharing of derived alleles between a series of West Eurasians (*X*) and population *Y* with respect to Ust’-Ishim (Fig. 2). Admixture between *X* and *Y* is expected to lead to an increase in the proportion of shared derived alleles. We set *Y* in our analyses to East, West and Central Africans and Oceanians, in which the value of the *D* statistic should not be significantly different from 0 if there is no admixture between *X* and *Y*.

### Testing for the presence of Basal Eurasian ancestry

We used the statistic *f_4_(West Eurasian W, East Asian or early hunter gatherer X; Ust’-Ishim, Chimp)* to look for the presence of Basal Eurasian ancestry in a West Eurasian *W* (Fig. S1) (19). This statistic tests if the data is consistent with a tree in which *W* and *X* lineages form a clade, which results in *f_4_* statistic not significantly different from 0. Significantly negative values are evidence for an affinity between the *W* and *X* lineages, which has been most parsimoniously explained by *W* carrying some ancestry from a population that split off from the non-African lineage prior to the separation of Ust’-Ishim (19).

### Simulations of selection

We used the simulation framework SLiM 2 (31) to build a realistic model of the human genome with empirical distributions of functional regions and selection coefficients, extending and generalizing a strategy previously applied by Harris and Nielsen (6). To obtain the positions of regions under negative selection, we downloaded coordinates of different classes of functionally important genomic regions from the Ensembl database (Ensembl Genes 91 and Ensembl Regulation 91), and converted them to BED format (32). In each simulation, we encoded those regions in a genomic structure in SLiM’s Eidos programming language, maintaining the distances between them. In order to model the heterogeneity of recombination rate along a genome in our simulations, we used empirically estimated genetic distances between all simulated genomic features using a recombination map inferred by the HapMap project (http://hapmap.ncbi.nlm.nih.gov/downloads/recombination/2011-01_phaseII_B37/) (33). To approximate a distribution of fitness effects (DFE) of new deleterious mutations, we used the DFE estimated from the frequency spectrum of human non-synonymous mutations, which represents a mixture of strongly, weakly and nearly-neutral mutations (34). The rate of accumulation of new mutations was set to 1×10^−8^ per bp per generation. The simulations themselves were performed in two steps (Fig. 4A), using a combination of human and Neandertal demographic models used in previous introgression studies (4, 6). In the first step, we simulated a simplified demography of modern humans and Neandertals prior to the introgression, starting with a burn-in period of 70,000 generations, to let the simulated genomes with mutations reach an equilibrium state (the length of this burn-in period was determined as 7 * ancestral human Ne, which was therefore set to a constant 10,000). The split of Neandertals and modern humans was set to 500,000 years ago, with *N_e_* of Neandertals and modern humans set to constant values of 1,000 and 10,000, respectively. This burn-in period was performed for each specific simulation scenario separately. At the end of the burn-in step, we simulated the split of African and non-African populations at 55 thousand years ago. Following the split, the non-African population experiences a bottleneck with *N_e_* = 1861 (as inferred by Gravel et al. (35)). All simulated individuals and accumulated mutations were saved to a population output file for use in the second step.

In the second step, we simulated a single pulse of admixture from Neandertals into the non-African population at a rate of 10%. To track Neandertal ancestry along simulated genomes through time, we placed 50,000 *neutral Neandertal markers* outside of any potentially functional sequence (which was determined as a union of all annotated Ensembl regions mentioned above) (Fig. 4A). The locations of these markers were randomly sampled from the set of nearly-fixed Yoruba-Neandertal differences present on the archaic admixture array (SNP panel 4 in (24)). Furthermore, to be able to track Neandertal ancestry within regions directly under negative selection, we placed additional set of fixed Neandertal markers within those regions (Fig. 4A).

Because the efficacy of selection is related to the *N_e_* of the population under consideration (36) we evaluated different demographic models for non-Africans, including a widely-used model by Gravel et al. (35) (i.e. long bottleneck followed by a period of exponential growth), a model of initial slow linear growth post admixture, as well as a model of constant *N_e_* after Neandertal introgression (Fig. S8). However, because we found that the *N_e_* of the admixed non-African population did not have an impact on the slope of the trajectory of Neandertal ancestry over time (Fig. S4), the main results in our paper were performed using a demographic model with constant *N_e_* = 10,000.

To track dynamics of selection over time, we periodically saved the simulation state, saving all mutations still segregating at each time-point (both neutral markers and deleterious modern human and Neandertal mutations) in a sample of 500 individuals in VCF format for further analysis. For efficiency reasons only simulation states in generations 1-10, 20, 50, 100 and then every 200th generation until the final generation 2200 (i.e. 55 thousand years, assuming generation time of 25 years) were saved.

### Evaluating the effect of negative selection against introgression

All of the following analyses were performed on VCF outputs from 20 replicates of our SLiM simulations, described in the previous paragraph. Trajectories of Neandertal ancestry in a population over time (Fig. 4B) were calculated by averaging the frequencies of all *neutral Neandertal markers* in a simulation in each time point across 500 sampled individuals. Analysis of the efficacy of selection against introgression as a function of distance from regions carrying deleterious variants (Fig. 4C) was performed by binning the 50,000 *neutral Neandertal markers* into 5 quintiles, based on their distance from the nearest region under selection. The lowest bin “0” contains *neutral Neandertal markers* within regions that carried accumulated variants. Neandertal ancestry proportions were then calculated for each of the 1,000 sampled chromosomes in each bin, combined from all simulation replicates. Analysis of allele frequency changes over time was performed by calculating the frequency change of each class of mutation (neutral Neandertal markers within and outside of selected regions, and deleterious mutations) between each consecutive pair of sampled time-points, and then averaged over all mutations. For example, if *x* and *y* are allele frequencies of a mutation at time-points *a* and *b*, then the allele frequency change was calculated as *(x – y) / (a – b)*. This calculation was repeated for all 20 simulation replicates and mean frequency changes were plotted for each replicate separately (Fig. 4D).

### Simulations of gene-flow between non-Africans and Africans

We simulated different scenarios of gene-flow between Africans and non-Africans after Neandertal introgression using the neutral coalescent programming framework msprime (37) (Fig. S2). We used the following demographic parameters: split of a chimpanzee lineage at 6 million years ago, split of Neandertals from anatomically modern humans at 500 kya, a split within Africa at 150 kya, and split of non-Africans from one of the two African lineages at 60 kya with a 5 ky long bottleneck of *N_e_* = 2000. We simulated a single 3% pulse of Neandertal admixture into a constant-size non-African population (*N_e_* = 10,000) at 55 kya. We sampled one chimpanzee chromosome, 4 Neandertal chromosomes sampled at 80 kya, single chromosomes from the non-African lineage sampled at time-points corresponding to dates of Upper-Paleolithic samples from our data, two sets of 50 chromosomes from the two present-day African populations and a set of 20 present-day non-African chromosomes. We simulated 500 Mb chromosomes using a mutation rate of 1×10^−8^ mutations per bp per generation and a recombination rate of 1×10^−8^ crossovers per bp per generation. We converted the sampled chromosomes into a table of all simulated SNPs (to represent genome-wide capture data similar to the set of 2.2 million sites from SNP panels 1, 2, 3 from (24)). For some analyses, we also generated a second set of SNPs by filtering only for sites carrying fixed African-Neandertal differences (to approximate the ascertainment of the archaic admixture array – SNP panel 4 in (24)). To estimate the true Neandertal ancestry levels we examined the origin of each simulated mutation in msprime and extracted only those SNPs that truly originated in the Neandertal population. Using this set of sites avoids any issues caused by introduction of Neandertal alleles into Africans via gene-flow from admixed non-Africans.

We evaluated the behavior of the admixture array ancestry proportion and direct and indirect *f_4_*-ratio estimates under three scenarios: (i) no gene-flow between Africans and non-Africans post Neandertal admixture, (ii) gene-flow from non-Africans into both African populations starting at 20 kya, (iii) gene-flow from one African population into non-Africans starting at 20 kya, (iv) bi-directional gene-flow between Africans and non-Africans starting at 20 kya, (v) increasing the total amount of functional sequence (thereby increasing the number of accumulated deleterious variants in Neandertals). In all scenarios of gene-flow, we used a moderate level of admixture of 0.0001 migrants per generation, which falls within the range of previously estimates levels of gene-flow between both populations (35, 38). The time of 20 kya was chosen to approximately match the results of our *D* statistic-based analysis of the affinity between Europeans and Africans over time (Fig. 2).

Using the simulated SNP sets (all SNPs and archaic admixture array-like set), we calculated direct and indirect f_4_-ratio estimates, as well as admixture array proportion estimates, as described above. Unbiased levels of Neandertal ancestry were calculated on the set of true Neandertal-derived SNPs. As the statistics can be relatively noisy, we simulated 100 replicates of each gene-flow scenario, and calculated average values of each individual statistic.

## Acknowledgements

We wish to thank Montgomery Slatkin, Benjamin Peter and Fabrizio Mafessoni for helpful discussions and comments on the manuscript. The study was funded by the Max Planck Society and the European Research Council (grant agreement number 694707 to S.P.)

---

**Figure S1.**
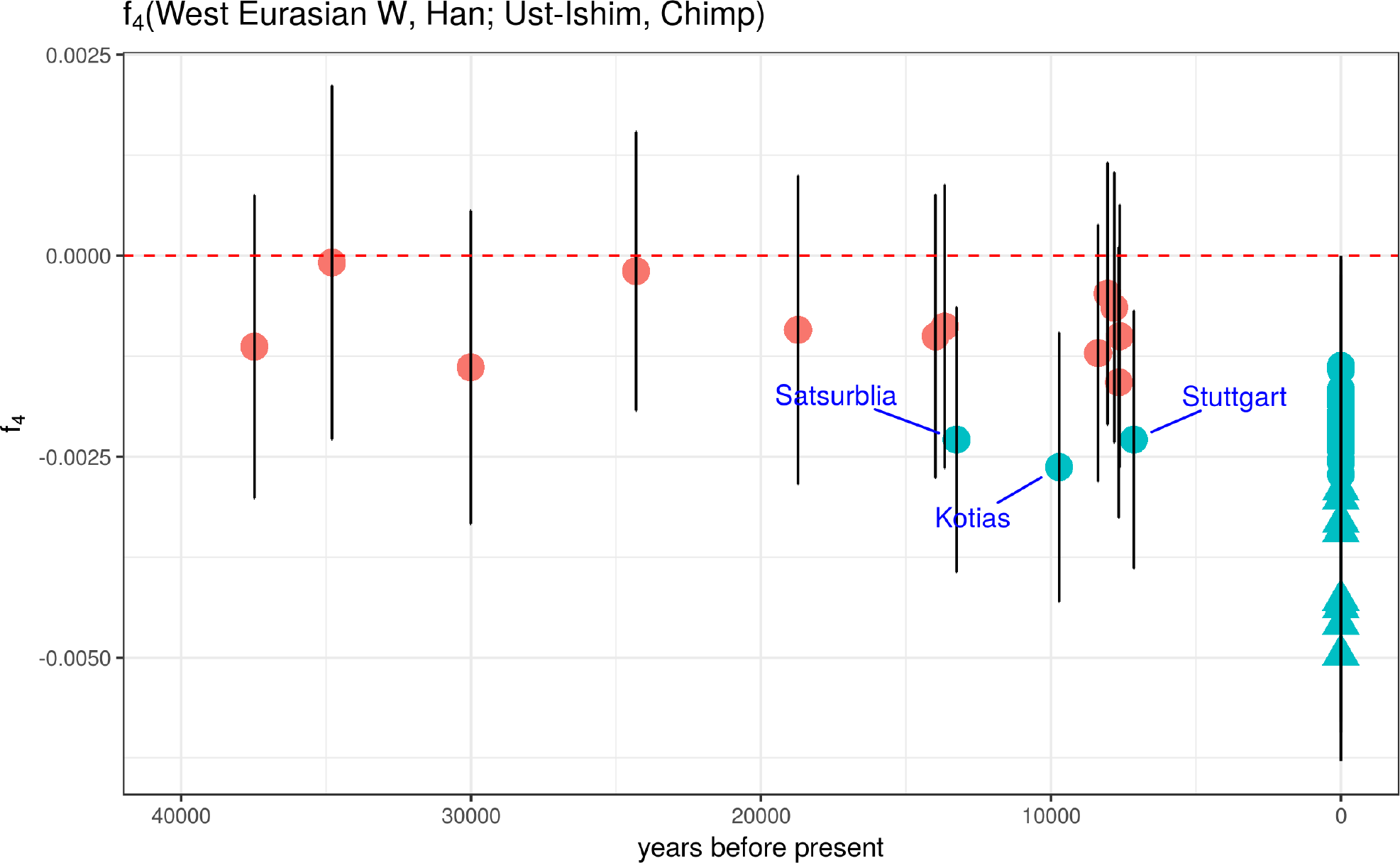
A signal of Basal Eurasian ancestry in West Eurasia over time. The statistic *f_4_(West Eurasian W, East Asian or ancient hunter gatherer X; Ust’-Ishim, Chimp)* has been previously used as a test of the presence of Basal Eurasian ancestry in a West Eurasian *W* (19). Specifically, it tests whether a population tree in which W and X lineages form a clade is consistent with the observed data, which results in *f_4_* statistic ~0. On the other hand, significantly negative values are evidence for an affinity of East Asian and Ust’-Ishim lineages, which can be most parsimoniously explained by *W* carrying an ancestry component from a population that split off from the non-African lineage prior to the separation of Ust’-Ishim. This “ghost” population is commonly referred to as Basal Eurasians (19). By analyzing a combined early-modern and present-day West Eurasian dataset, we find that this *f_4_* statistic becomes consistently negative in the present, which is in agreement with the hypothesis that present-day West Eurasians carry (in different proportions) Basal Eurasian ancestry that was not present in early European hunter gatherers. Figure shows samples with at least 500k captured sites. The three highlighted ancient Europeans are those who have been confirmed previously to carry a statistically significant amount of Basal Eurasian ancestry. Present-day individuals are Europeans (circles) and Near Easterners (triangles) from the SGDP panel (29).

**Figure S2.**
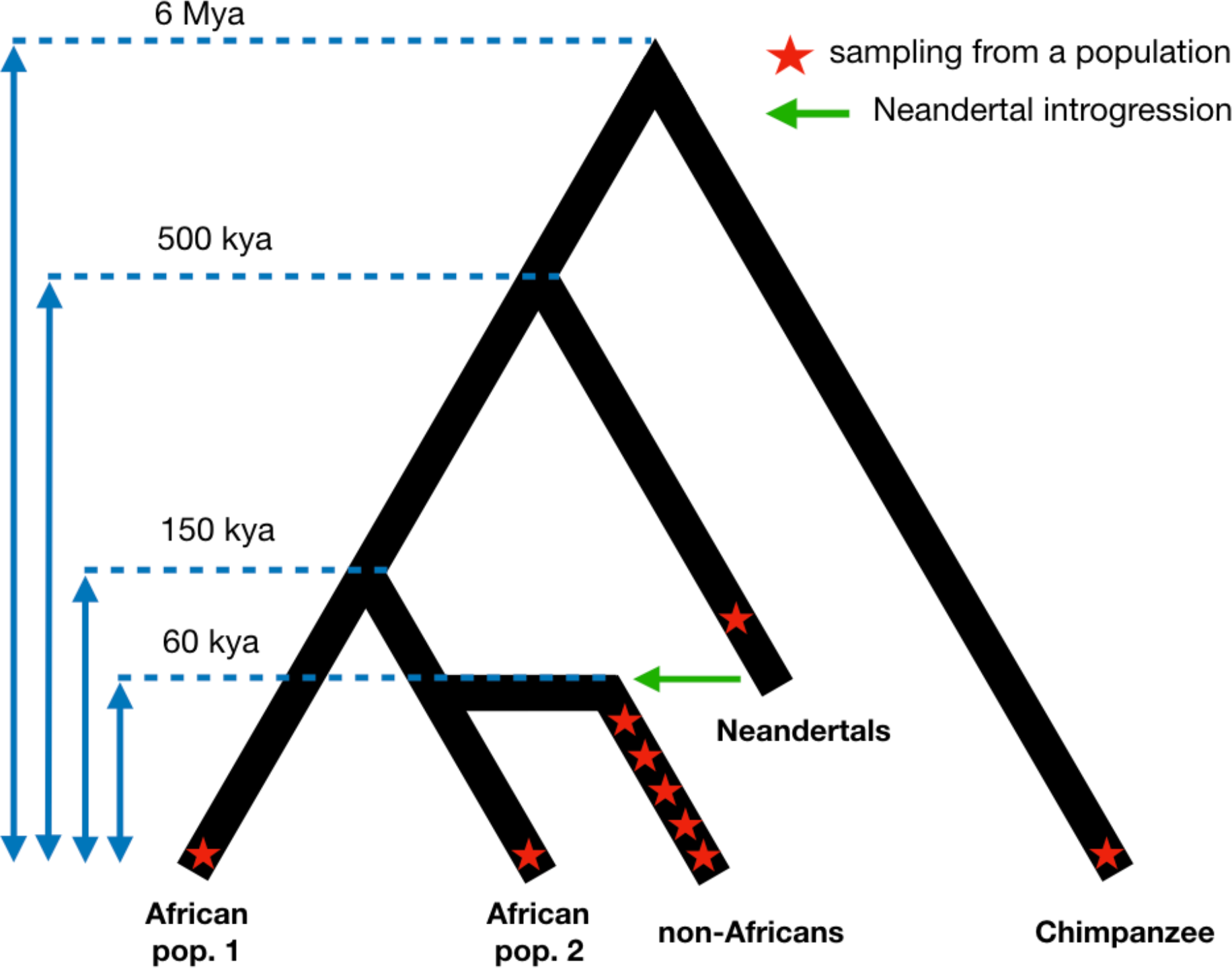
Demographic model used for testing the robustness of admixture statistics. Blue dashed lines show split times between simulated populations, red stars indicate approximate points in time at which simulated 500 Mb chromosomes were sampled.

**Figure S3.**
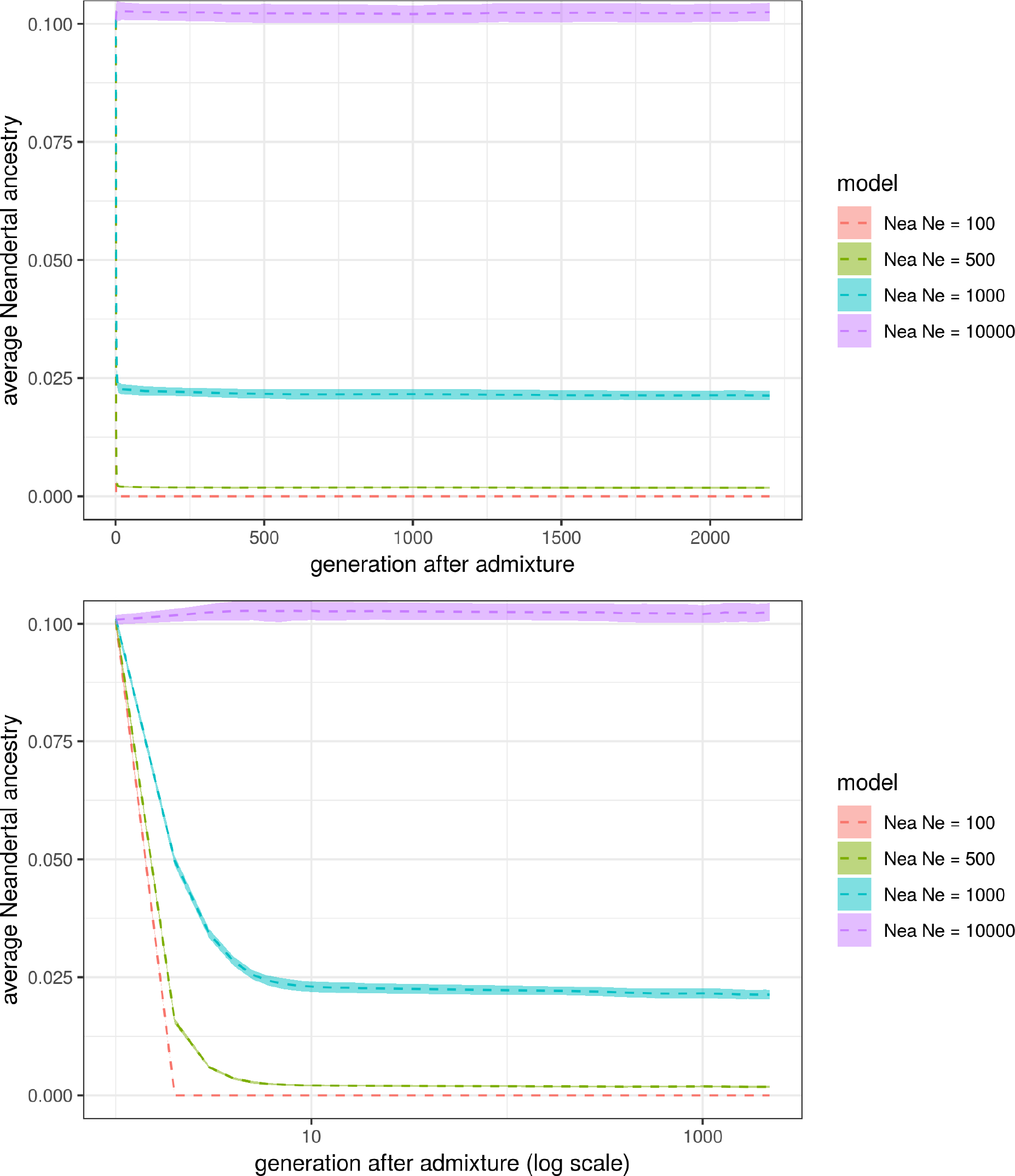
The effect of Neandertal *N_e_* (Nea *Ne*) on trajectories of Neandertal ancestry after introgression. Upper and lower panels show linear and logarithmic timescales, respectively. The lower the *N_e_* of Neandertal population, the more deleterious alleles behave nearly neutrally, allowing them to reach high frequencies in the Neandertals (6, 7). This imposes a stronger genetic load of the initial modern-human-Neandertal hybrids, causing a more abrupt removal of Neandertal ancestry in the generations shortly after admixture.

**Figure S4.**
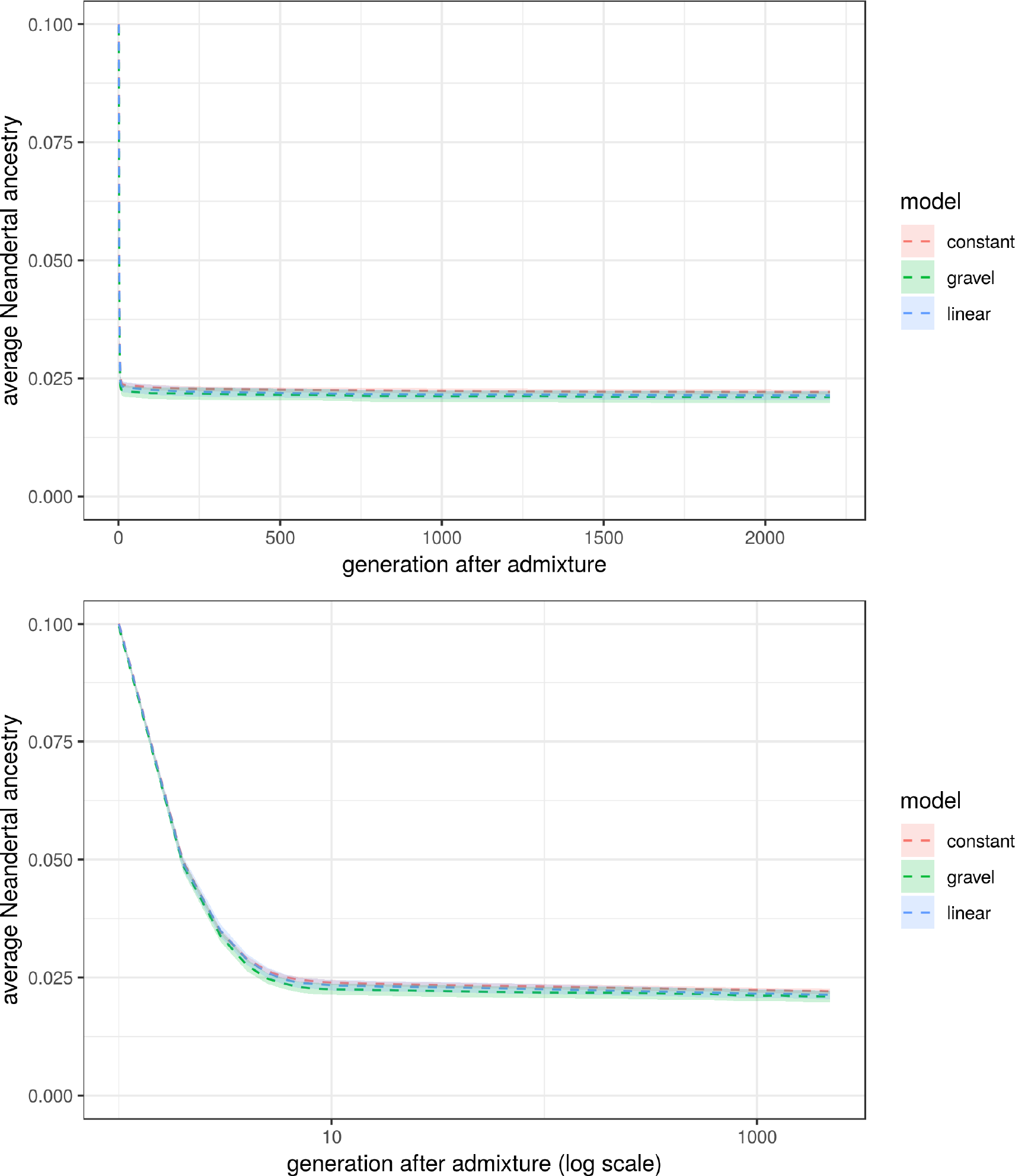
The effect of non-African demography on trajectories of Neandertal ancestry after introgression. Upper and lower panels show linear and logarithmic timescales, respectively. Although *N_e_* as a function of time differs dramatically between all three demographic models that we considered (Fig. S8), changing this parameter does not have a strong impact on the overall shape of Neandertal ancestry trajectories.

**Figure S5.**
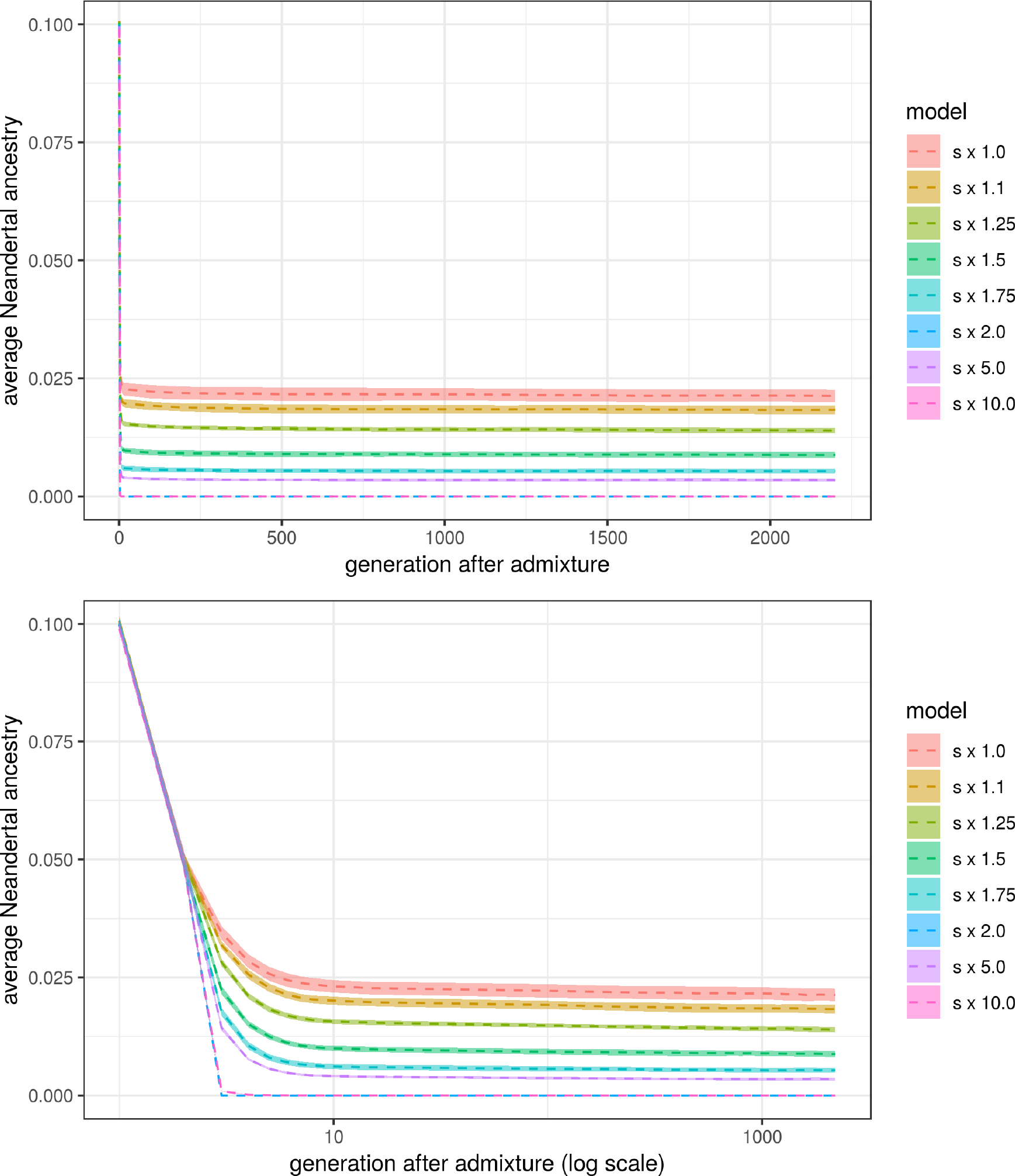
The effect of making Neandertal mutations more deleterious by increasing their selection coefficients. Upper and lower panels show linear and logarithmic timescales, respectively. We artificially increased the selection coefficient *s* of introgressed Neandertal alleles by multiplying their *s* by a constant factor. We find that this affects only the final level of Neandertal ancestry in the population, due to stronger genetic burden on hybrids in the first generations after admixture.

**Figure S6.**
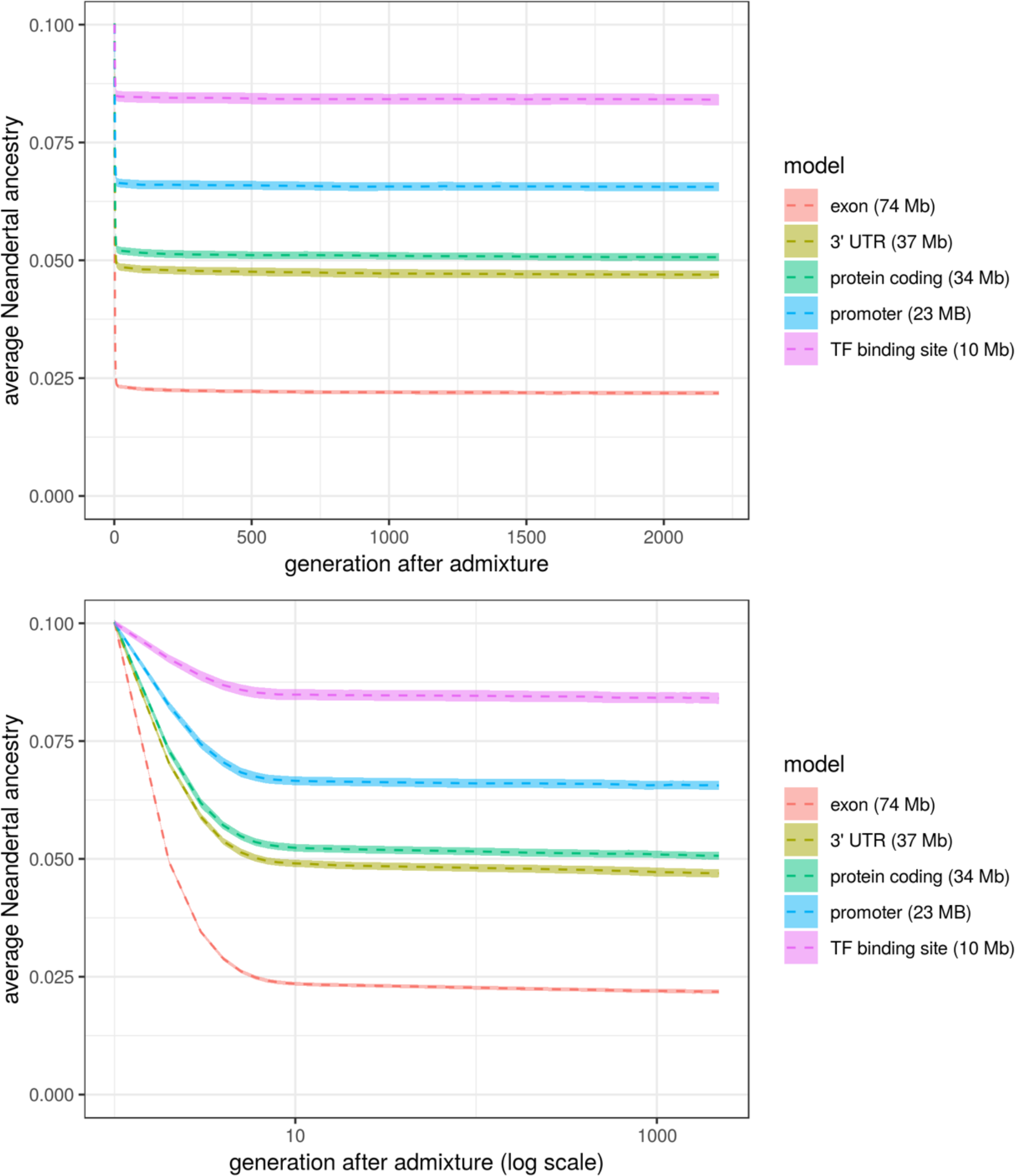
The effect of changing the total amount of potentially deleterious sequence. Upper and bottom panels show linear and logarithmic timescales, respectively. We simulated deleterious mutations in either full exonic, 3’ UTR, protein coding, promoter, or TF binding site regions. Simulations with larger “targets” for deleterious mutations have lower final levels of Neandertal ancestry.

**Figure S7.**
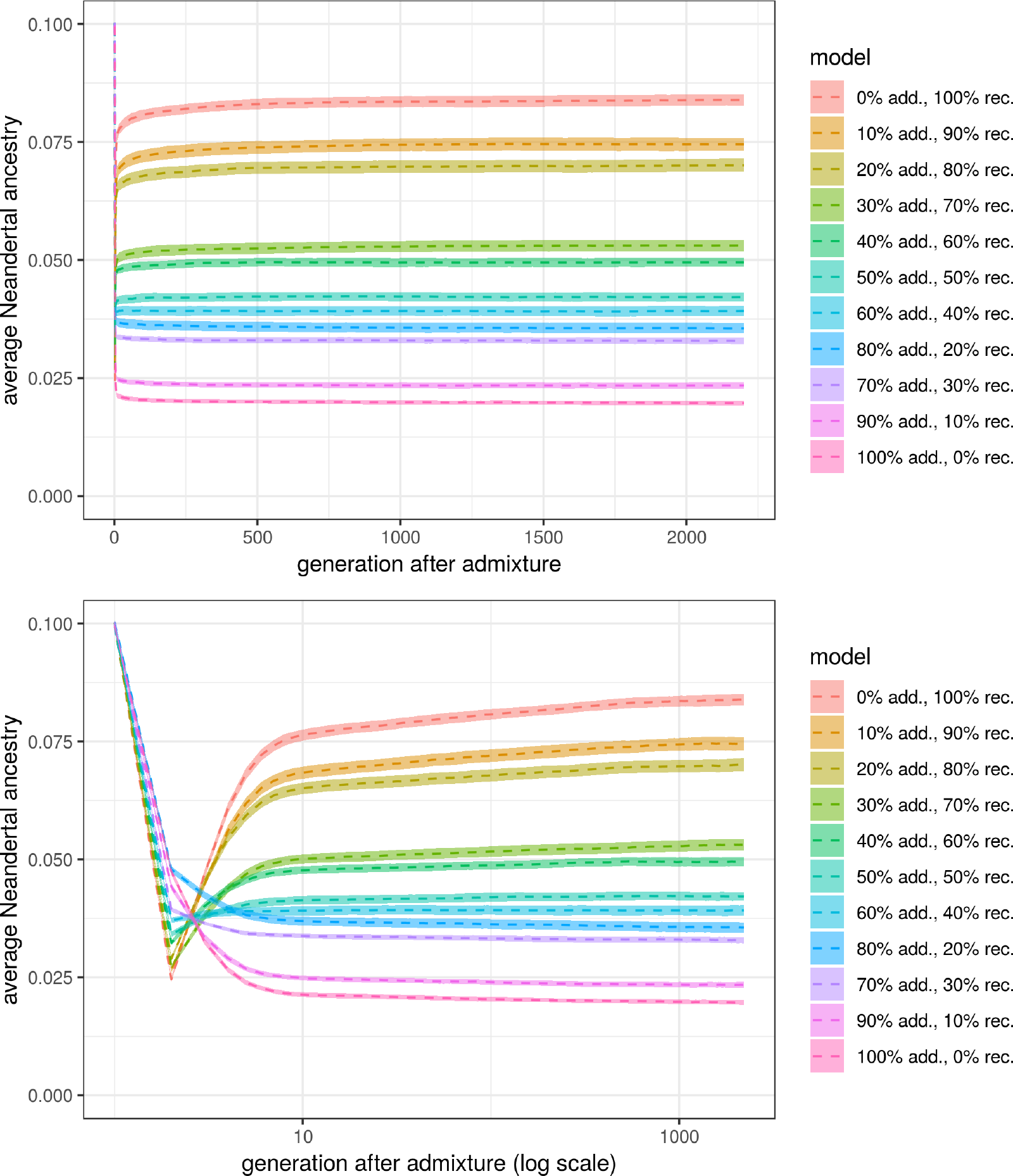
The effect of changing the proportions of recessive and additive mutations. Upper and lower panels show linear and logarithmic timescales, respectively. It has been shown that the dominance coefficient of deleterious mutations can lead to Neandertal ancestry trajectories following entirely opposite patterns (6). Specifically, models with only recessive mutations lead to an initial increase of the Neandertal ancestry proportions due to heterosis (6). Due to these opposing effects of dominance, we investigated scenarios with different mixtures of dominance coefficients of deleterious mutations. We found that changing the ratios of recessive and additive mutations affects only the final baseline of Neandertal ancestry in the population, and does not lead to a steady decline in Neandertal ancestry over time.

**Figure S8.**
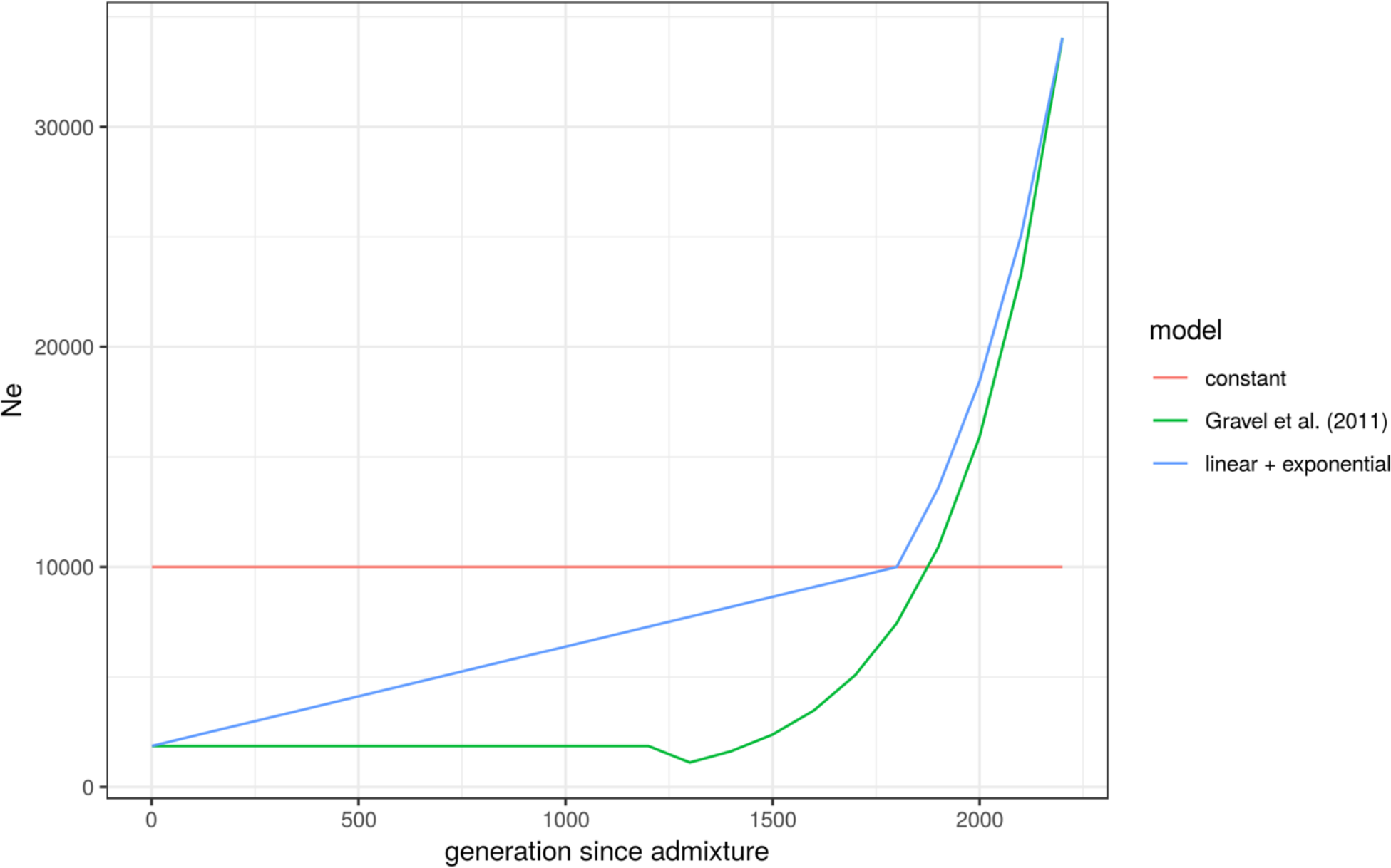
Three models of non-African demography after Neandertal admixture. *Ne* as a function of time for three models of non-African demography: a model of constant *Ne* after Neandertal introgression, a model of initial slow linear growth post admixture, and a long bottleneck followed by a period of exponential growth (Gravel et al. (35)). Unless otherwise noted, all analyses in this paper use the constant *N_e_* model.

**Figure S9.**
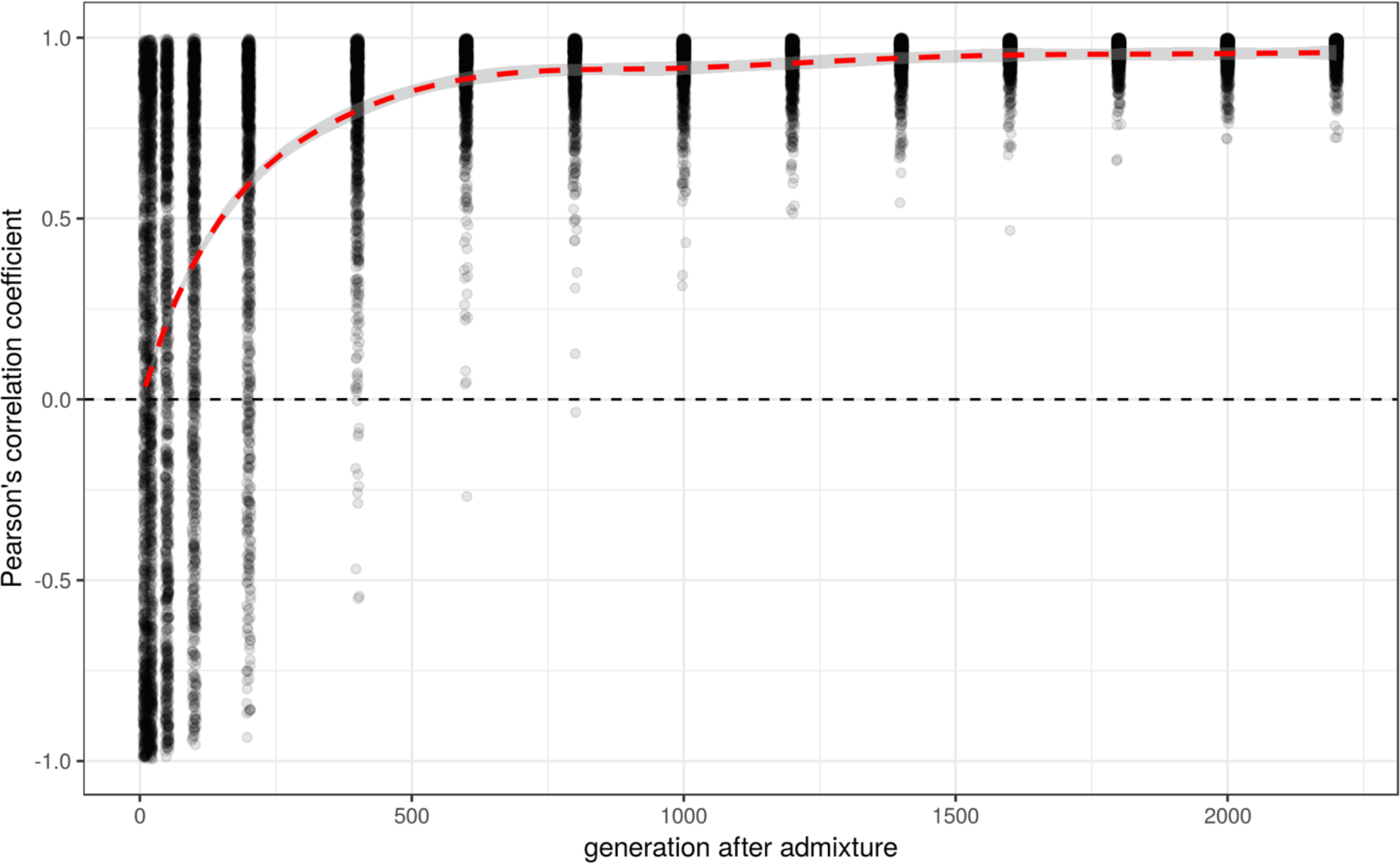
Coefficients of correlation between the proportion of surviving Neandertal ancestry and distance to a genomic region under negative selection. Pearson’s correlation coefficient of a correlation between Neandertal ancestry proportion and the distance to the nearest functional region, at a given point in time. Each dot represents a single simulated “individual” with 500,000 informative sites. This figure uses the same data presented in Fig. 4C (which shows averages over all simulations at each individual time-point).

## References

1. Green RE, et al. (2010) A Draft Sequence of the Neandertal Genome. Science 328(5979):710–722.

2. Sankararaman S, Patterson N, Li H, Pääbo S, Reich D (2012) The Date of Interbreeding between Neandertals and Modern Humans. PLOS Genet 8(10):e1002947.

3. Sankararaman S, et al. (2014) The genomic landscape of Neanderthal ancestry in present-day humans. Nature 507(7492):354–357.

4. Vernot B, Akey JM (2014) Resurrecting Surviving Neandertal Lineages from Modern Human Genomes. Science 343(6174):1017–1021.

5. Schumer M, et al. (2018) Natural selection interacts with recombination to shape the evolution of hybrid genomes. Science:eaar3684.

6. Harris K, Nielsen R (2016) The Genetic Cost of Neanderthal Introgression. Genetics:genetics.116.186890.

7. Juric I, Aeschbacher S, Coop G (2016) The Strength of Selection against Neanderthal Introgression. PLOS Genet 12(11):e1006340.

8. Fu Q, et al. (2016) The genetic history of Ice Age Europe. Nature 534(7606):200–205.

9. Harris K, Nielsen R (2017) Q&A: Where did the Neanderthals go? BMC Biol 15:73.

10. Yang MA, Fu Q (2018) Insights into Modern Human Prehistory Using Ancient Genomes. Trends Genet 0(0). doi:10.1016/j.tig.2017.11.008.

11. Reich D (2018) Who We Are and How We Got Here: Ancient DNA and the New Science of the Human Past (Pantheon, New York).

12. Steinrücken M, Spence JP, Kamm JA, Wieczorek E, Song YS Model-based detection and analysis of introgressed Neanderthal ancestry in modern humans. Mol Ecol 0(0). doi:10.1111/mec.14565.

13. Prüfer K, et al. (2017) A high-coverage Neandertal genome from Vindija Cave in Croatia. Science:eaao1887.

14. Racimo F, Sankararaman S, Nielsen R, Huerta-Sánchez E (2015) Evidence for archaic adaptive introgression in humans. Nat Rev Genet 16(6):359–371.

15. Patterson N, et al. (2012) Ancient Admixture in Human History. Genetics 192(3):1065–1093.

16. Peter BM (2016) Admixture, Population Structure and F-Statistics. Genetics:genetics.115.183913.

17. Prüfer K, et al. (2014) The complete genome sequence of a Neanderthal from the Altai Mountains. Nature 505(7481):43–49.

18. Hajdinjak M, et al. (2018) Reconstructing the genetic history of late Neanderthals. Nature 555(7698):652–656.

19. Lazaridis I, et al. (2016) Genomic insights into the origin of farming in the ancient Near East. Nature 536(7617):419–424.

20. Vernot B, Akey JM (2015) Complex History of Admixture between Modern Humans and Neandertals. Am J Hum Genet 96(3):448–453.

21. Lazaridis I, et al. (2014) Ancient human genomes suggest three ancestral populations for present-day Europeans. Nature 513(7518):409–413.

22. Schlebusch CM, et al. (2017) Southern African ancient genomes estimate modern human divergence to 350,000 to 260,000 years ago. Science:eaao6266.

23. Skoglund P, et al. (2017) Reconstructing Prehistoric African Population Structure. Cell 171(1):59–71.e21.

24. Fu Q, et al. (2015) An early modern human from Romania with a recent Neanderthal ancestor. Nature 524(7564):216–219.

25. Sankararaman S, Mallick S, Patterson N, Reich D (2016) The Combined Landscape of Denisovan and Neanderthal Ancestry in Present-Day Humans. Curr Biol 26(9):1241–1247.

26. Jacobsen F, Omland KE Increasing evidence of the role of gene flow in animal evolution: hybrid speciation in the yellow-rumped warbler complex. Mol Ecol 20(11):2236–2239.

27. Cui R, et al. Phylogenomics Reveals Extensive Reticulate Evolution in Xiphophorus Fishes. Evolution 67(8):2166–2179.

28. Schrider DR, Ayroles J, Matute DR, Kern AD (2018) Supervised machine learning reveals introgressed loci in the genomes of Drosophila simulans and D. sechellia. PLOS Genet 14(4):e1007341.

29. Mallick S, et al. (2016) The Simons Genome Diversity Project: 300 genomes from 142 diverse populations. Nature 538(7624):201–206.

30. Meyer M, et al. (2012) A High-Coverage Genome Sequence from an Archaic Denisovan Individual. Science 338(6104):222–226.

31. Haller BC, Messer PW (2017) SLiM 2: Flexible, Interactive Forward Genetic Simulations. Mol Biol Evol 34(1):230–240.

32. Zerbino DR, et al. (2018) Ensembl 2018. Nucleic Acids Res 46(D1):D754–D761.

33. International HapMap Consortium, et al. (2007) A second generation human haplotype map of over 3.1 million SNPs. Nature 449(7164):851–861.

34. Eyre-Walker A, Woolfit M, Phelps T (2006) The Distribution of Fitness Effects of New Deleterious Amino Acid Mutations in Humans. Genetics 173(2):891–900.

35. Gravel S, et al. (2011) Demographic history and rare allele sharing among human populations. Proc Natl Acad Sci 108(29):11983–11988.

36. Lanfear R, Kokko H, Eyre-Walker A (2014) Population size and the rate of evolution. Trends Ecol Evol 29(1):33–41.

37. Kelleher J, Etheridge AM, McVean G (2016) Efficient Coalescent Simulation and Genealogical Analysis for Large Sample Sizes. PLOS Comput Biol 12(5):e1004842.

38. Harris K, Nielsen R (2013) Inferring Demographic History from a Spectrum of Shared Haplotype Lengths. PLOS Genet 9(6):e1003521.

